# The SEA-AD DREAM Challenge: Community benchmarking human and AI agent solutions for Alzheimer’s disease neuropathology prediction from single-nucleus transcriptomics

**DOI:** 10.64898/2026.07.02.736180

**Authors:** Hsin-Yu Lai, Nikolaos Kalavros, Verena Chung, Eitan S. Kaplan, Julio Saez-Rodriguez, Lun Ai, Dimitris Anastassiou, Lingyi Cai, Erica Chen, Ignacio Garach Vélez, Gamze Gürsoy, Luis Javier Herrera, Xiaoting Li, Eric Londin, Phillipe Loher, Iliza Nazeraj, Francisco Ortuño, Tai-Hsien Ou Yang, Isidore Rigoutsos, Ignacio Rojas, The SEA-AD DREAM Community, Gaia Andreoletti, Luca Foschini, Laura Heath, Tomiko Oskotsky, Marina Sirota, Gustavo Stolovitzky, Kyle J. Travaglini, James Zou, Mariano I. Gabitto

## Abstract

Single-nucleus transcriptomic atlases offer an unprecedented opportunity to connect cellular molecular states with Alzheimer’s disease (AD) neuropathology, but whether these profiles encode reproducible, predictive information about pathological burden remains unclear. We present the SEA-AD DREAM Challenge, an open, international, model-to-data competition built on the Seattle Alzheimer’s Disease Brain Cell Atlas to predict Alzheimer’s disease neuropathological severity from single-nucleus RNA-sequencing data. Participants developed containerized models to predict categorical neuropathological staging, including overall Alzheimer’s disease neuropathologic change, Braak stage, Thal phase, and CERAD score, as well as quantitative amyloid-β and phospho-tau burden measured by 6E10 and AT8 immunohistochemistry. Across 17 eligible teams from 15 countries, the crowdsourcing framework enabled systematic comparison of diverse computational approaches and surfaced a broad landscape of modeling strategies and candidate predictive features. Top-performing methods achieved near-perfect prediction of categorical staging, with the best submission reaching a quadratic weighted kappa of 1.0 for the Overall AD Neuropathological Change score (ADNC), and competitive prediction of quantitative pathological burden in held-out data, with a best concordance correlation coefficient of 0.48. Post hoc perturbation analyses revealed that top categorical-stage predictions relied heavily on donor-level metadata-driven signals rather than transcriptomic features, whereas quantitative pathology prediction was more robust and supported by transcriptomic and cell-type-associated features with potential biological relevance to AD progression. The challenge also introduced the first AI Agent Track in a DREAM Challenge, providing an early benchmark for autonomous and human-guided agentic model development in single-cell neuroscience. This work demonstrates that single-nucleus transcriptomes encode substantial information about Alzheimer’s disease pathology, establishes a reproducible benchmark for molecular neuropathology prediction, and highlights critical principles for designing privacy-preserving, leakage-aware community challenges using deeply phenotyped human brain data.

## Introduction

Alzheimer’s disease (AD) is the most common neurodegenerative disorder, affecting approximately 1 in 9 individuals (10.9%) aged 65 and older and leading to progressive cognitive decline, dementia, and substantial personal, social, and economic burden on patients and their families [1]. At the neuropathological level, AD is defined by two hallmark lesions: extracellular amyloid-beta (Aβ) plaques and intraneuronal aggregates of hyperphosphorylated tau (neurofibrillary tangles, NFTs) distributed across partially overlapping neuroanatomical axes [2–5]. Staging systems that quantify the anatomical spread of these lesions like Thal phases for Aβ [3], Braak stages for NFTs [4], and CERAD scores for neuritic plaque density [5], are integrated into the NIA-AA neuropathological framework that defines Overall AD Neuropathological Change (ADNC) [2]. These staging systems remain the gold standard for post-mortem diagnosis and serve as essential endpoints for both clinical-pathologic correlation studies and the development of antemortem biomarkers. Critically, neuropathological staging is available for many cohorts where matched molecular profiling remains limited, restricting our ability to systematically link transcriptomic signatures to disease severity.

Single-nucleus RNA sequencing (snRNA-seq) enables transcriptomic profiling of post-mortem human brain tissue with cell-type resolution, generating datasets of extraordinary richness and dimensionality [6,7]. These datasets have begun to reveal molecular signatures of cellular vulnerability and resilience in aging and neurodegeneration. However, most available cohorts lack paired high-resolution quantitative neuropathological measurements, restricting analyses to primarily case-versus-control comparisons or broad histopathological disease stages [8–12]. A smaller subset of studies have paired single-cell data with regional or per-cell pathological burden quantification [13,14], demonstrating that cell-type-specific transcriptomic changes track closely with the accumulation of amyloid and tau pathology. These observations raise a fundamental question: Do single-nucleus transcriptomic profiles contain reproducible signals associated with Alzheimer’s disease neuropathological burden? Moreover, can these signals support cross-region prediction of amyloid and tau pathology? If transcriptomic states contain sufficient information about neuropathological burden, it should be possible to train computational models to mechanistically understand the molecular underpinnings of neuropathological measures through the lens of gene expression. Evaluating the extent and limits of this predictive capacity has the potential to provide insight into cell types and gene programs that encode neuropathology progression. In addition, if accurate prediction is possible, it will enable molecular neuropathology at scale in cohorts lacking direct histopathological assessment.

The Seattle Alzheimer’s Disease Brain Cell Atlas (SEA-AD) consortium, a collaboration among the Allen Institute for Brain Science, the University of Washington, and the Adult Changes in Thought (ACT) study at Kaiser Permanente Washington Health Research Institute, has generated a uniquely comprehensive, openly accessible resource for the AD research community [15]. The SEA-AD dataset integrates deep single-nucleus multi-omics profiling with quantitative neuropathology and rich clinical phenotyping across a cohort of 84 deeply characterized post-mortem brain donors spanning the full spectrum of AD severity.

Neuropathological staging (ADNC, Braak, Thal, CERAD, LATE-NC [16], Lewy Body Disease stage [17]) was performed by the University of Washington BRaIN Laboratory, and quantitative immunohistochemical burden for six protein markers (6E10, AT8, NeuN, GFAP, pTDP43, aSyn) was derived from whole-slide image analysis using HALO software on sections immediately adjacent to those used for snRNA-seq.

DREAM (Dialogue for Reverse Engineering Assessments and Methods) Challenges are open, international crowdsourced competitions that leverage the collective intelligence of the research community to address problems that no single group can solve alone [18,19]. First proposed in 2006, DREAM Challenges have repeatedly demonstrated that assembling diverse community solutions can outperform even the best individual models, an effect termed the “wisdom of crowds,” across domains ranging from gene regulatory network inference [20] and drug sensitivity prediction [21] to microbiome science [22] and clinical prediction using electronic health records [23]. Crucially, this performance advantage derives not only from raw model diversity but also from the structured, unbiased evaluation enabled by withholding test labels until final scoring.

Our hypothesis is that predicting AD neuropathological burden from single-cell transcriptomics is precisely the kind of high-dimensional, multi-faceted problems for which crowdsourcing approaches can excel. No consensus methodological framework exists; the feature space spans tens of thousands of genes measured across hundreds of thousands of nuclei, ground-truth labels span ordinal, semi-continuous, and continuous scales, and the biological relationships between transcriptomic state and pathological burden are expected to be non-linear and potentially cell-type-specific. Engaging the global community ensures exposure to the full spectrum of approaches and the resulting diversity of solutions may generate biological insights beyond what performance rankings alone convey.

In this work, we present the SEA-AD DREAM Challenge, an open, community-driven benchmarking competition for predicting Alzheimer’s disease neuropathological burden from single-nucleus transcriptomic data. By combining a well-phenotyped transcriptomic dataset with a model-to-data evaluation framework, the challenge enables rigorous and privacy-preserving comparison of diverse computational approaches. Across two complementary tasks: predicting donor-level neuropathological staging and quantitative pathological burden, we systematically assess the extent to which transcriptomic signals, alone or in tandem with patient-level categorical data, encode disease severity. The resulting collection of models provides both a performance benchmark and a resource for identifying transcriptomic features and modeling strategies that are most informative for neuropathology prediction.

## Results

### Overview

Here we describe the design and execution of the SEA-AD DREAM Challenge, an open international crowdsourced competition to predict quantitative AD neuropathological burden from single-nucleus RNA-sequencing (snRNA-seq) data derived from the SEA-AD cohort. The challenge comprised two complementary prediction tasks. Task 1 asked participants to predict categorical neuropathological severity at the donor level: the primary endpoint was the overall AD Neuropathological Change (ADNC) summary score [2], with secondary endpoints of Braak neurofibrillary tangle stage [3], Thal amyloid phase [4], and CERAD neuritic plaque score [5]. Task 2 asked participants to predict continuous immunohistochemistry-quantified pathological burden for two canonical AD markers: 6E10 (pan-amyloid) and AT8 (phospho-tau Ser202/Thr205), using quantitative image analysis scores derived from the SEA-AD neuropathological pipeline [15]. An exploratory AI Agent Track, the first agentic track incorporated into a DREAM Challenge, additionally invited LLM-based agents to compete on both tasks with limited human-in-the-loop model development (where humans can offer feedback as an advisor but AI agents should be the main contributor), providing an early benchmark for agentic computational biology in comparison to the human track. Prediction performance was evaluated using quadratic weighted kappa (QWK) [24] for Task 1 and the concordance correlation coefficient (CCC) [25] for Task 2 (averaged over 6E10 and AT8). QWK assesses agreement between predicted and true ordinal labels while penalizing larger discrepancies more heavily, whereas CCC evaluates agreement between continuous predictions and observations by jointly measuring correlation (precision) and deviation from perfect concordance (accuracy).

In parallel with the challenge, the organizing team applied scVIP [26], an interpretable generative model linking single-cell transcriptomics to donor-level phenotypes, as a complementary organizer-side model to contextualize community submissions; scVIP was trained on both SEA-AD and ROSMAP data [27, 28] and was not subject to the challenge submission constraints (see Methods).

### The DREAM Challenge Data

We provided training data consisting of snRNA-seq profiles from the middle temporal gyrus (MTG) and dorsal frontal cortex (DFC) of the full SEA-AD cohort (n = 84 donors), both of which have been previously publicly released. The validation and test datasets were derived from the superior temporal gyrus (STG) and inferior temporal gyrus (ITG), respectively, using a subset of these donors without major co-morbidities (n = 43 donors for each set) (see Methods section about the training and validation/testing data for more details). The corresponding quantitative neuropathology measurements for these regions have not been publicly released. Participants interacted with the validation and test datasets through a Docker-based evaluation environment (see Methods). In addition to the snRNA-seq profiles, we provided accompanying metadata including anonymized donor ID, sex, age at death, race, years of education, and post-mortem interval, as these variables (with the exception of donor ID) may provide additional biological or clinical context relevant to AD progression. Although the donors in the validation and test datasets were a subset of those in the training data, donor identifiers were reassigned so that donor IDs could not be directly matched across dataset splits. Cell type annotations were also provided (annotation procedure described in Figure 1 and in [15] for more detail), as selective cellular vulnerability is known to play an important role in AD progression. Through this challenge, we aimed to establish an open and reproducible benchmark for transcriptomic prediction of AD neuropathology and demonstrate the value of crowdsourcing for accelerating methodological progress in single-cell neuroscience.

**Figure 1:**
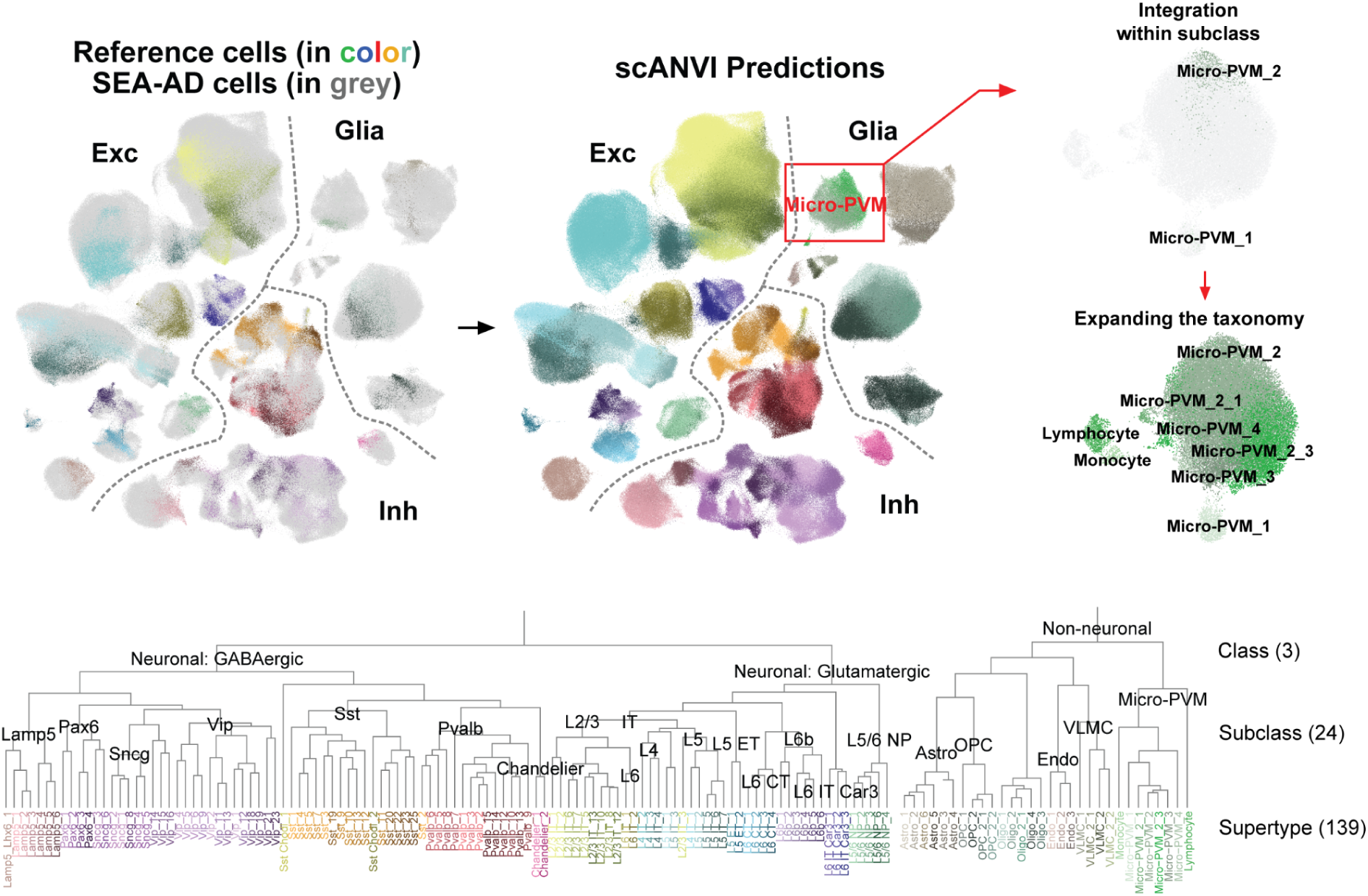
(Top). Data from SEA-AD (grey) was mapped to a BICAN reference taxonomy (in color). Clusters of cells that were out of the reference cell distribution were added to the taxonomy for non-neuronal/glial subclasses (i.e., Astrocytes, Oligodendrocytes, Oligodendrocyte Precursor Cells (OPCs), Microglia/Perivascular Macrophages/Other Immune cells (Micro-PVM), Endothelial Cells, and Vascular Leptomeningeal Cells and Fibroblasts (VLMCs). (Bottom). Dendrogram showing the relationship between the three levels of the taxonomy that includes Class (3 types), Subclass (24 types), and Supertype (139 types).

### The DREAM Challenge Results

A total of 190 Synapse users from 15 countries registered for the Challenge (see Supplementary Figure 1), of whom 17 teams submitted at least one eligible scored model. The detailed numbers of teams and submissions for the validation and the test rounds of the challenge are shown in Supplementary Table 1. Below, we summarize the main results.

#### Categorical staging prediction achieved high apparent performance but was sensitive to donor-associated shortcuts

In Task 1, we evaluated ADNC prediction using quadratic weighted kappa (QWK). To account for stochastic variability, we performed 10,000 bootstrap re-samplings of cells from the test dataset and re-evaluated all methods, excluding the complementary organizer-side model, scVIP, where only 5 bootstraps were performed due to computational cost. Because the training, validation, and test datasets share the same donors across different brain regions, and donor-level metadata are available, participants could have inferred mappings between donors across splits. Since ADNC is constant per donor, such mappings could yield artificially perfect predictions.

Consistent with this possibility, team gisl7 achieved perfect accuracy across all bootstrap samples (Figure 2), suggesting substantial reliance on donor-identifying signals (see Data Leakage Analysis below). In contrast, teams CMC-TJU and BioICAR achieved near-perfect performance, with median bootstrapped QWK values of approximately 0.98 and 0.93, respectively, suggesting that their predictions may rely less on these signals. The complementary organizer-side model, scVIP, does not use metadata but trained on both ROSMAP and SEA-AD data, achieved QWK values around 0.65, demonstrating that transcriptomic signals alone still contain substantial information predictive of AD staging.

**Figure 2.**
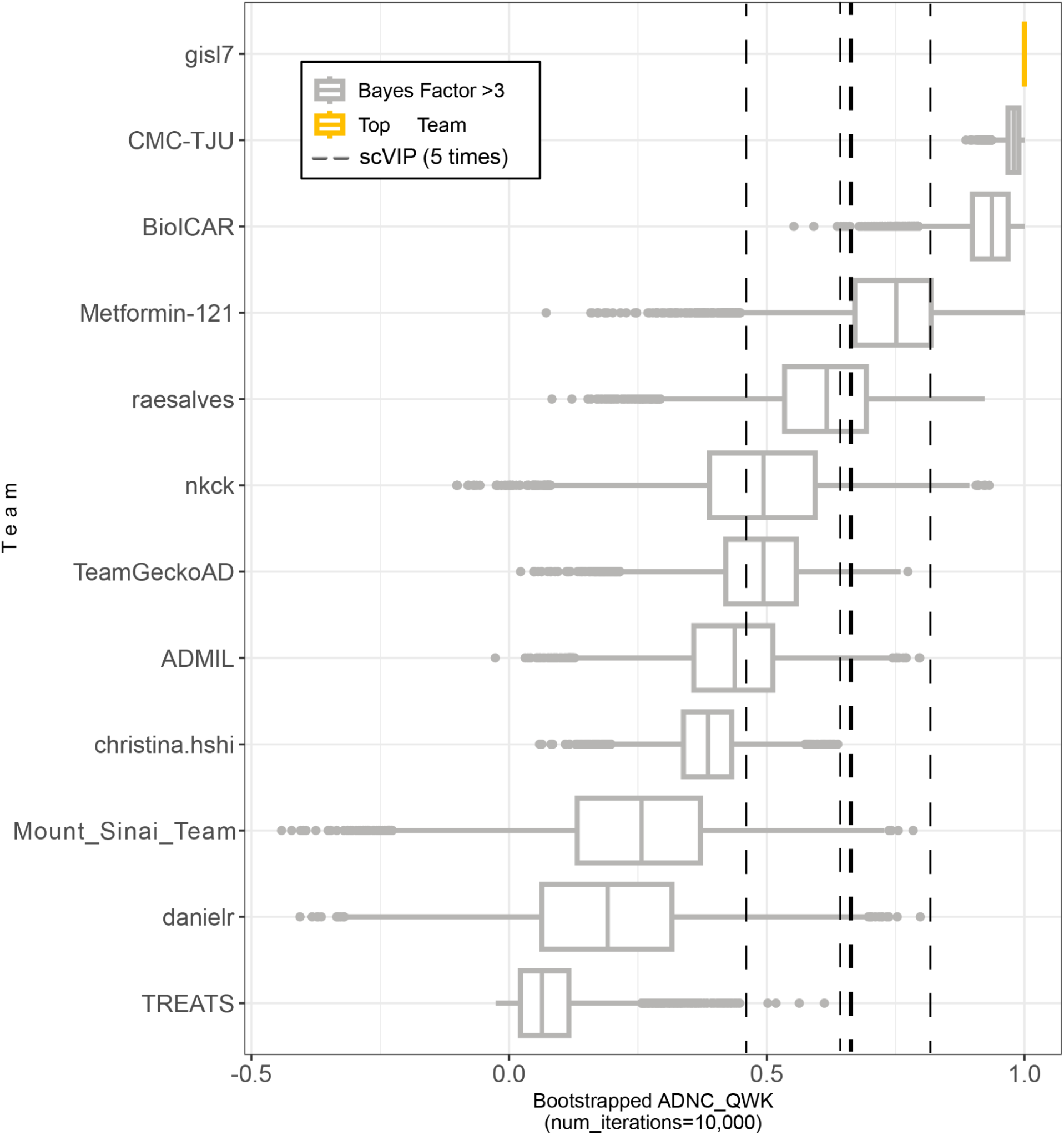
Bootstrapped ADNC prediction performance (10,000 iterations) for each team that submitted a model to the Final Round, compared with five bootstrap runs of scVIP, the complementary organizer-side model.

To further assess the contribution of metadata-driven leakage, we evaluated model performance under randomized metadata settings. Specifically, we performed three complementary tests. In Test 1, metadata and labels were jointly shuffled across donors; under this setting, models relying primarily on metadata should retain much of their performance. In Test 2, only metadata were shuffled while labels were preserved; models that do not depend on metadata should remain largely unaffected. In Test 3, the order of cells (i.e., the rows of the anndata object) was shuffled to assess whether any methods implicitly relied on donor or cell ordering. All teams were evaluated by comparing their official leaderboard performance with their performance under these perturbation settings. To estimate variability in the top-performing methods, the analyses were repeated 10 times for the top three teams on each leaderboard. The same evaluation framework was then applied to Task 2 submissions.

For the top-performing teams in Task 1 (Figure 3), we observed distinct patterns of dependence on metadata and transcriptomic features. The performance of gisl7 remained stable under Test 1 but degraded substantially under Test 2, consistent with strong reliance on metadata-driven signals. In contrast, the model from team CMC-TJU showed performance degradation under both Test 1 and Test 2, suggesting that it integrates both metadata and cell- or gene-level features. For team BioICAR, performance degraded only under Test 1, indicating little or no direct reliance on metadata. These interpretations are consistent with independent inspection of the submitted code.

**Figure 3.**
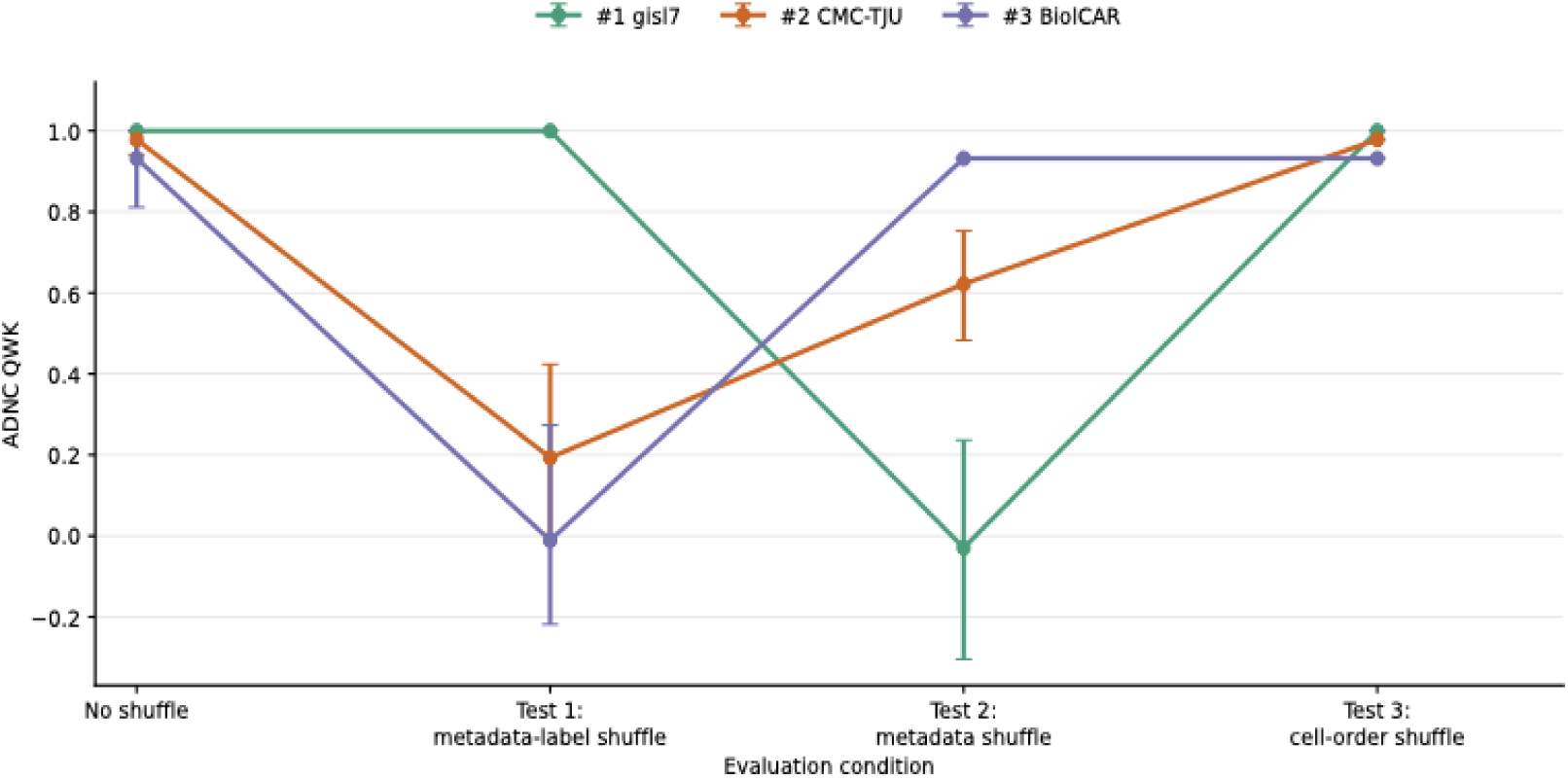
Performance of the top three teams on (i) the official leaderboard in Task 1, (ii-iv) under the data leakage test, where (ii) in test 1, the metadata and the labels were shuffled together; (iii) in test 2, the metadata were shuffled; and (iv) in test 3, the order of the cells was shuffled.

#### Quantitative regional pathology prediction was more difficult and less completely explained by donor metadata

In Task 2, we evaluated the prediction of amyloid-β and tau burden using the mean concordance correlation coefficient (CCC) across 6E10 and AT8. As in Task 1, we performed 10,000 bootstrap re-samplings and re-evaluated all methods, excluding scVIP, for which only five bootstrap runs were performed due to computational cost. In contrast to Task 1, no model achieved perfect performance (Figure 4), likely reflecting the fact that quantitative pathological burden varies across brain regions within individual donors. Team CMC-TJU achieved a median bootstrapped CCC of approximately 0.48, while the top nine teams and scVIP all achieved CCC values greater than 0.25. The organizer-side model, scVIP, achieved a CCC of approximately 0.53, comparable to the top-performing methods. These results suggest that Task 2 is less completely explained by donor-identity-based shortcuts and is more likely to depend on transcriptomic and cellular signals associated with regional pathology.

**Figure 4.**
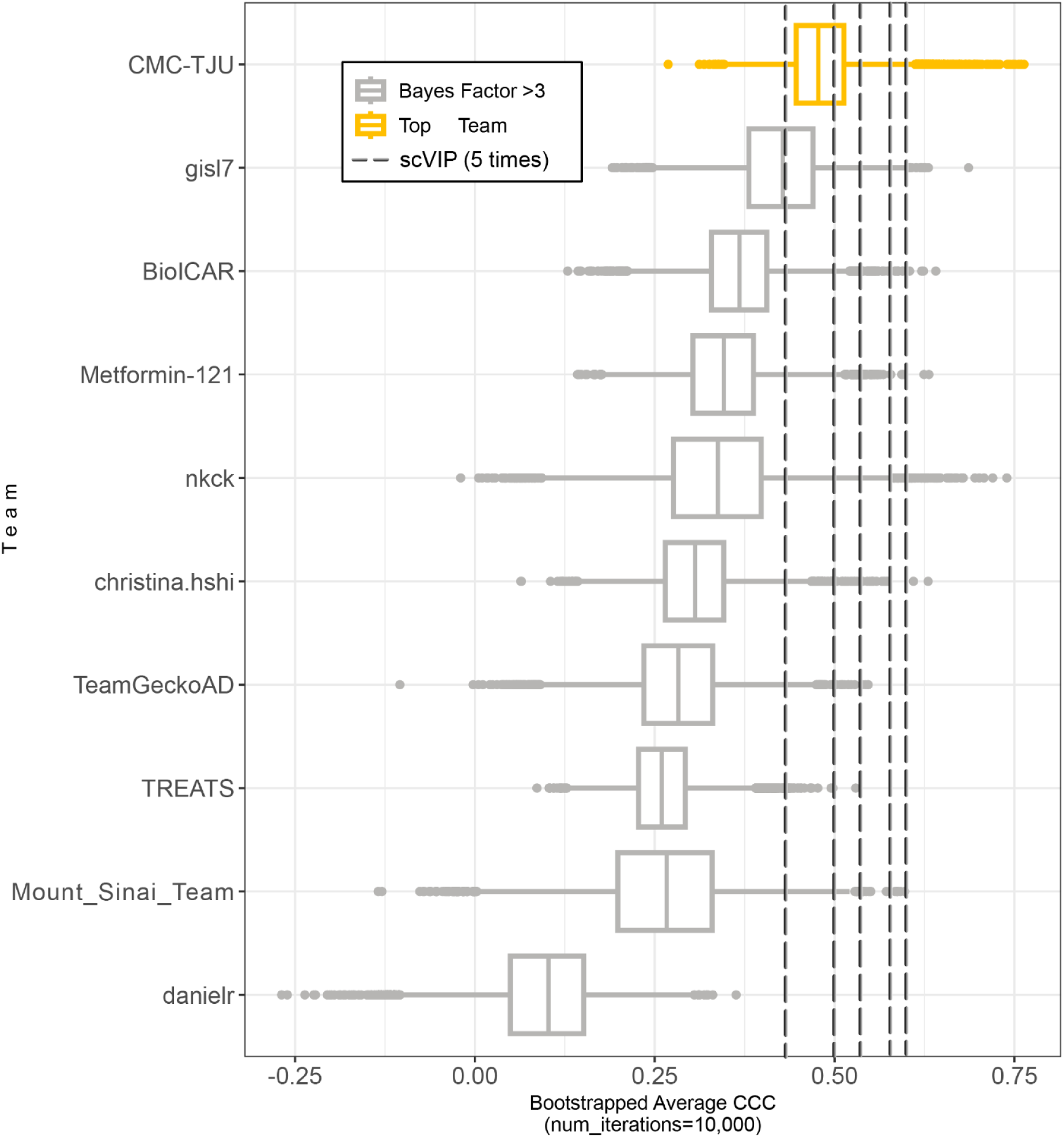
Bootstrapped 6e10 and AT8 prediction performance (10,000 iterations) for each team that submitted a model to the Final Round, compared with five bootstrap runs of scVIP, the complementary organizer-side model.

We next applied the same perturbation framework to Task 2 submissions. In contrast to Task 1, the perturbation results did not support a simple donor-matching explanation for top Task 2 performance. As shown in Figure 5, both gisl7 and CMC-TJU exhibited performance patterns consistent with mixed use of donor-level metadata and transcriptomic features, whereas BioICAR again showed no evidence of metadata dependence. Across both tasks, Test 3 (cell-order shuffling) had no measurable effect on model performance, suggesting that none of the submitted methods exploited cell-ordering artifacts.

**Figure 5.**
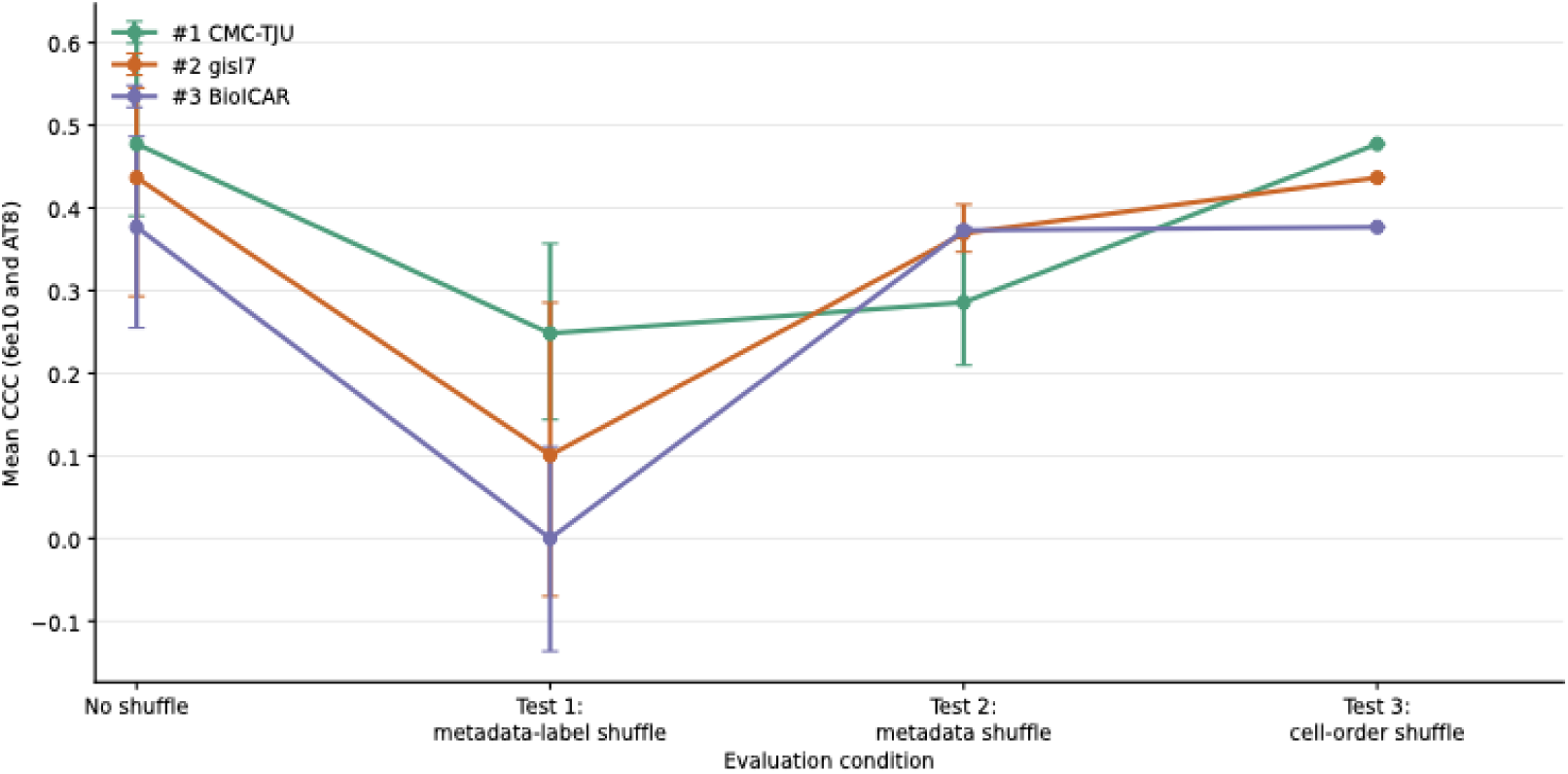
Task 2 performance of the top three teams on (i) the official leaderboard and (ii-iv) under the data leakage test, where (ii) in test 1, the metadata and the labels were shuffled together; (iii) in test 2, the metadata were shuffled; and (iv) in test 3, the order of the cells was shuffled.

#### Top-performing approaches used cell-type-aware donor representations and identified diverse features

We next examined the modeling strategies and the feature-importance patterns of the top-performing teams (team BioICAR, CMC-TJU, and gisl7; more information can be found in Methods). The three top-performing teams used distinct approaches, but all preserved some form of cellular context before donor-level prediction.

Team BioICAR developed a two-stage modeling framework that first generated predictions from each cell supertype (as defined in [15]) and then integrated these predictions to produce donor-level outputs. Rather than modeling individual cells directly, the method summarized each donor–supertype pair by averaging gene expression across cells of the same supertype. These donor–supertype profiles were then compressed using principal component analysis, and separate models were trained for each cell supertype. In the second stage, the cell-supertype-level predictions were concatenated and used as input to a final donor-level model. Because the first-stage models were predominantly linear, predictive features could be examined through model coefficients and mapped back to gene-level contributions through PCA back-projection.

Team CMC-TJU used a more restricted feature set and trained predictors at the level of broad cellular classes (3 classes) rather than individual cell supertypes (139 supertypes). The method limited the input space to 3,430 nervous system genes enriched for primate- and human-specific sequence motifs (“pyknons”) [29] and grouped cells into three broad cellular classes: GABAergic neurons, glutamatergic neurons, and non-neuronal cells. In actuality, only 3,157 of the 3,430 genes were included in the training data and used in these analyses. To increase the number of training observations, cells within each donor–class pair were divided into 100 equal subgroups, each summarized by mean expression. Donor metadata, including sex, years of education, and APOE genotype, were appended to each subgroup-level expression profile, and separate LASSO regression models were trained for each outcome variable.

Team gisl7 summarized each donor and brain region using gene-level expression fractions, defined as the fraction of cells with nonzero expression for each gene. The team additionally constructed gene-pair co-expression features from the microglia–perivascular macrophage subclass, selected because of its relevance to Alzheimer’s disease pathology and consistent representation across brain regions. After choosing the expression-fraction and Micro-PVM co-expression features using mutual information, these features and donor-level covariates were combined in supervised regression models trained separately for each pathology target.

To further understand the molecular and metadata-derived features prioritized by these models, we performed feature-importance analyses (see Methods; Table 1 for the top five features per task and team, and Supplementary Table 2 for the top 100 Task 2 features per team). The identified features varied across teams, consistent with differences in model architecture, feature construction, and the cellular resolution at which predictors were defined. The highest-ranked features included both well-characterized AD-related signals and genes with less evident connections to AD. The latter may represent understudied biology, correlated proxies for disease-associated cellular states, or unstable associations arising from the high dimensionality of the feature space relative to the cohort size.

**Table 1.** Top features selected by each model.

| Team | Task | Identified features |
| --- | --- | --- |
| BiolCAR | Task 1 | Supertypes: Micro-PVM_1, VLMC_1, Vip_14, Sst_4<br>Genes: *ADAMTS9-AS2*, *NRG1*, *GRID2*, *CNTN5* |
|  | Task 2 | AT8:<br>Supertypes: Monocyte, OPC_1, SMC-SEAAD, and Micro-PVM_1<br>Genes: AL592183.1, ARL17B, TOGARAM2, CENPP, AC105402.3<br>6e10:<br>Supertypes: Sst_3, OPC_1, L4 IT_2, and L5/6 NP_1<br>Genes: AC105402.3, NUTM2B-AS1, AC119673.2, MT-CO1, AC068587.4<br>Metadata: PMI, Race (choice=White), Sex, Years of education, Age at Death, APOE Genotype |
| CMC-TJU | Task 1 | Genes: *OR12D3*, *NBPF4*, *TFAP2B*<br>Metadata: years of education, sex, and APOE genotype |
|  | Task 2 | 6e10:<br>Genes: *OR12D3*, *LBX1*<br>Metadata: APOE genotype<br>AT8:<br>Genes: *OR12D3*, *HOXB8*, *NBPF*<br>Metadata: APOE genotype |
| gisI7 | Task 2 | 6e10:<br>Genes: AC016590.1, HLA-DRB5, AC026341.2, AC009654.1, TBC1D3, SHISA6<br>Co-expressions within Micro-PVM: PTPRG-MSR1, IPCEF1-CPM, OXR1-RGL1<br>Metadata: APOE 4/4, APOE 3/4, Age at Death<br>AT8:<br>Genes: AC025887.2, PCED1B, AC010201.3, PKN2-AS1, AC114296.1, MKRN3<br>Co-expressions within Micro-PVM: PTPRG-SAMD4A, PTPRG-MSR1, PTPRG-MYO1E, PTPRG-MITF<br>Metadata: APOE 4/4 |

When the analysis was expanded to the top 100 features per team, broader collections of AD-relevant genes and cell populations became apparent. BioICAR’s supertype-level features encompassed Micro-PVM, OPC, Sst, Vip, and excitatory neuronal populations. Associated genes included *DLGAP2* and *CHL1*, which are involved in neuronal and synaptic organization; *SPARCL1*, an astrocyte- and vascular-associated extracellular-matrix protein with synapse-modulating functions; *SCN1A* and *NOVA1*, which contribute to inhibitory-neuron function and neuronal RNA regulation; and *ATG10* and *LIPA*, which participate in autophagy and lysosomal lipid metabolism. *WWOX*, located at a replicated late-onset AD GWAS locus and reported to interact with tau, was also among the highly ranked features.

CMC-TJU’s features included genes related to inflammatory and neurotrophic signaling, including *CHI3L2* and *NTRK1*. *CHI3L2* has been associated with inflammatory responses and AD-related molecular changes, although it is less established as an AD cerebrospinal-fluid biomarker than the related protein CHI3L1/YKL-40. *NTRK1* encodes TrkA, the high-affinity NGF receptor required for trophic support of basal-forebrain cholinergic neurons.

The expanded microglial feature set from gisl7 included inflammatory-signaling genes such as *IL1RAP* and *IRAK3* and calcium-signaling genes including *CACNA1D* and *ITPR1*. *HLA-DRB5* lies within the AD-associated *HLA-DRB5/DRB1* region, although the specific causal gene or variant within this locus remains unresolved. *MARK3*, another highly ranked feature, belongs to the MARK kinase family and has been linked to phosphorylation of tau at Ser262, but evidence connecting this event to early AD pathology is stronger for *MARK4* and the temporal position of Ser262 phosphorylation remains unsettled.

These findings indicate that the submitted models prioritized features spanning microglial inflammatory signaling, neuronal and synaptic biology, neurotrophic pathways, lysosomal and autophagic processes, AD-associated genetic regions, and established donor-level risk factors such as APOE genotype. These associations are consistent with the multicellular and multifactorial pathology of Alzheimer’s disease, although feature importance should be interpreted as predictive association rather than evidence of causal involvement.

#### The first DREAM Agent Track revealed feasibility and current limits of agentic model development

Two teams submitted to the AI Agent Track. The first (Nguyen & Zhan, New Mexico Tech) constructed a pipeline combining library-size normalization, dimensionality reduction via 3,000-component PCA, per-endpoint MLP classifiers for ordinal targets, and Random Forest regressors for continuous targets, with donor-level aggregation; the authors explicitly describe this submission as a human-controlled execution of LLM-generated code rather than an autonomous agent, with humans retaining responsibility for debugging, scaling corrections, and architecture pivots. The second (UWisc-Madison) implemented a multi-round optimization multi-agent AutoML framework [30] with specialized agents for instruction parsing, data planning, model selection, and code execution, performing iterative refinement over successive rounds and converging on an ordinal XGBoost ensemble over PCA-embedded, cell-type-aggregated donor features.

On Task 1, the first submission achieved ADNC QWK = 1.00 (Braak 0.78, Thal 0.99, CERAD 1.00), matching the top-performing human team on the primary metric and suggesting that AI agents also learned to utilize the metadata information; the second achieved ADNC QWK = 0.53, near the median of human submissions. Only the first submission was scored for Task 2, where it achieved an average 6E10/AT8 CCC of 0.18 (6E10 0.27, AT8 0.10), below all but one human team (best 0.48; human range 0.11–0.48). Because the Agent Track was exploratory and not powered for formal statistical comparison, these results should be interpreted as descriptive; the Task 1 parity with the top human team reflects convergent exploitation of donor-level metadata or other dataset-level shortcuts rather than independent transcriptomic inference.

## Discussion

Through open crowdsourcing, the SEA-AD DREAM Challenge demonstrated that models trained on single-nucleus transcriptomic profiles from the middle temporal gyrus (MTG; Brodmann area 21) and dorsal frontal cortex (DFC; Brodmann area 9) can predict Alzheimer’s disease neuropathological burden in held-out superior temporal gyrus (STG) and inferior temporal gyrus (ITG) samples although the interpretation of predictive performance depends on potential leakage mechanism. Categorical donor-level staging achieved near-perfect apparent performance in some submissions but was sensitive to donor-associated shortcuts, whereas quantitative regional pathology prediction was more difficult and less completely explained by metadata or donor identity. Among 14 eligible final submissions for Task 1 and 11 for Task 2, top-performing teams achieved quadratic weighted kappa (QWK) scores of 1 for overall ADNC prediction and mean concordance correlation coefficients (CCC) of 0.48 for continuous immunofluorescence burden prediction.

Post hoc leakage analyses indicated that some submitted models leveraged correlations among donor metadata variables, including age, post-mortem interval, sex, and years of education, rather than relying solely on transcriptomic features. Team gisl7 showed a substantial performance decrease under these assessments, and Team CMC-TJU also showed reduced performance, suggesting partial reliance on metadata variables, which was later confirmed by reviewing the teams’ code and writeups. Team BioICAR represented a more nuanced case. Although their model did not explicitly use donor metadata, it relied on cell-type-stratified expression features; because cell-type abundances are highly correlated within donors across cortical regions [31], these features may have implicitly enabled approximate donor matching between training and test datasets. More broadly, because the challenge ranked teams based purely on predictive performance and included overlapping donors between training and test sets, it inadvertently rewarded signals that captured donor identity rather than transcriptomic disease signal alone. These findings highlight the importance of systematically evaluating potential leakage in future challenges involving richly phenotyped human cohorts. In contrast, Task 2 performance remained relatively robust under the leakage assessments, suggesting that top-performing models captured predictive signals beyond metadata or donor identity.

These results also offer broader methodological lessons for single-cell prediction tasks. Although the dataset contained a large number of cells, the effective sample size was limited by the 84 donors available for training, making this a high-dimensional, low-sample-size prediction problem at the donor level. The strongest submissions reflected this constraint: top-performing teams combined explicit feature engineering, careful feature selection, and relatively simple predictive models, suggesting that much of the neuropathology-associated signal can be captured through appropriately constructed donor-level representations. Notably, all three top-performing teams preserved some form of cellular context before donor-level prediction, including supertype-specific expression modeling by BioICAR, broad class-level modeling by CMC-TJU, and microglia–perivascular macrophage-specific co-expression analysis by gisl7. This convergence suggests that cell-type-aware representations may be useful for donor-level prediction from single-cell data, although the specific cell populations prioritized by these models were not always straightforward to interpret or directly aligned with previously reported SEA-AD disease-associated supertypes. In addition, most teams trained separate models for Task 1 and Task 2, limiting their ability to identify shared features underlying both neuropathological staging and quantitative pathology burden.

The organizer-side model, scVIP, was developed with a different and complementary objective. Unlike the top-performing submissions, which generally separated feature selection, representation construction, and downstream prediction, scVIP jointly learns cellular representations, cell-type-aware donor-level representations, and phenotype associations within a hierarchical generative framework. This end-to-end formulation provides a principled route to identifying vulnerable cell populations and associated gene programs, but it also introduces a larger parameter space and therefore greater data requirements. Accordingly, scVIP was trained jointly on ROSMAP and SEA-AD data to improve model stability and support cross-cohort analysis. As a result, the challenge submissions and scVIP highlight a tradeoff between data-efficient engineered representations optimized for within-cohort prediction and structured generative models designed for biological interpretation and cross-cohort generalization. Future work should evaluate these strategies under harmonized cross-cohort settings and develop methods that combine strong predictive performance with interpretable, cell-type-aware representation learning.

Feature-importance analyses across top-performing methods provided a complementary view of the molecular and metadata-derived signals used for prediction. Several prioritized features were consistent with AD-relevant biology, including APOE genotype, neuronal or synaptic genes, immune-associated features such as *HLA-DRB5*, and myeloid-associated predictors. At the same time, many selected genes were less canonical, noncoding, or difficult to interpret mechanistically. This heterogeneity highlights a key benefit of the crowdsourcing framework: by enabling diverse modeling strategies, the Challenge surfaced complementary predictive signals that can serve as hypotheses for future study. Because these signals were identified through predictive models rather than perturbational or causal analyses, their mechanistic significance remains to be established through replication, cross-cohort analyses, and targeted follow-up studies.

The AI Agent Track provided an early empirical data point on the capacity of autonomous agents for competitive model development leveraging single cell transcriptomics data. The two participating teams occupied opposite points on the human-in-the-loop spectrum: a human-orchestrated submission that used an LLM for code generation and a fully autonomous multi-agent framework. The human-orchestrated submission was more competitive on Task 1, consistent with the authors’ own report that domain judgment including choosing between RF and neural regressors, diagnosing silent 40× scaling errors, distinguishing real CCC gains from overfitting remained a human responsibility. The autonomous framework, while lower-scoring, successfully produced reproducible, containerized predictions without human intervention, demonstrating that end-to-end scientific autonomy is technically feasible even where it is not yet competitive. The Agent Track infrastructure, which includes standardized Docker environments, automated execution under identical resource constraints, fair scoring against identical held-out ground truth, and a structured AI-contribution checklist (hypothesis development, experimental design, data analysis, and observed limitations), provides a reusable template for future agentic benchmarks in computational biology.

More broadly, the SEA-AD DREAM Challenge illustrates several distinctive advantages of the crowdsourced challenge paradigm for neurodegeneration research. The participation of teams spanning computational biology, machine learning, neuroscience, statistics, and biomedical informatics brought a breadth of methodological approaches and a richer map of the solution landscape than any single group could have generated within a comparable timeline, leading to unexpected strategies that can inform subsequent hypothesis-driven work. This diversity was made possible in part by the model-to-data framework, which enabled rigorous, unbiased evaluation while protecting donor data privacy, a methodological standard that should be adopted more broadly for sensitive human brain datasets. Public leaderboards fostered transparency and rapid iteration, and the reproducibility requirements built into the challenge, including containerized submissions and mandatory code release, collectively constitute a reusable community resource for future researchers working with SEA-AD or analogous datasets, and establish a template for rigorous computational benchmarking in the neurogenomics community [32,33,34].

### Limitations

Several limitations of this Challenge should be acknowledged. First, the training, validation, and test datasets were derived from a single cohort (SEA-AD). While this design enabled consistent benchmarking, it led to reduced interpretability in Task 1 (due to shared AD staging across datasets) and limits the evaluation of the generalizability of top-performing models in Task 2 to other cohorts or tissue processing platforms.

Second, the generalizability of these findings to non-cortical regions may be limited. The training, validation, and test datasets all consist of cortical brain regions, with MTG included in the training data. In parallel with the Challenge, internal analyses revealed that, within cortical regions, cell-type abundances and molecular profiles are highly correlated across donors. The Challenge results are consistent with this observation, suggesting that such shared signals contribute substantially to predictive performance. However, these relationships may not extend to non-cortical brain regions, where cellular composition and transcriptional programs differ more substantially. As a result, while the Challenge reinforces the presence of strong, shared transcriptomic signals within cortical regions, the extent to which these models generalize beyond this context remains to be determined.

Third, the Challenge cohort, while deeply phenotyped, is of moderate sample size (84 donors total with 1-4 brain regions per donor), limiting statistical power to detect small differences in model performance and to characterize model behavior in demographic or genetic subgroups (e.g., stratified by APOE genotype or co-occurring pathologies). Fourth, the AI Agent Track was exploratory, with only two submissions (one scored for both Tasks, one scored for Task 1 only) and was not structured to enable formal statistical comparison with human teams; rigorous agentic benchmarking, including prespecified agent–human matchups, standardized compute budgets, and mandatory submission of agent thought-process and tool-use traces, is deferred to future Challenge iterations.

### Future directions

The SEA-AD DREAM Challenge highlights several high-priority directions for the field. A key consideration in this Challenge is the use of a single cohort, driven in part by the availability of well-characterized MTG and A9 datasets from SEA-AD. Given that generating high-quality transcriptomic data with matched neuropathological measurements is both resource-intensive and time-consuming, expanding such datasets remains a broader challenge for the field. This underscores the importance of aligning challenge design with data generation and release strategies at an early stage.

Looking ahead, extending prediction tasks to additional brain regions—particularly those spanning earlier and later stages of AD neuropathological progression, as well as non-cortical regions—would enable more comprehensive evaluation of model generalizability across the neuroanatomical cascade of disease. Multi-omic integration represents another important direction: incorporating chromatin accessibility (snATAC-seq), DNA methylation, or spatial transcriptomics alongside snRNA-seq may further improve both predictive performance and mechanistic interpretability. Finally, external validation of top-performing models on independent cohorts, as well as prospective evaluation in preclinical or mild cognitive impairment populations, will be essential for advancing these approaches toward clinical and translational applications.

The AI Agent Track raises exciting prospects for computational biology more broadly. As frontier LLMs improve in scientific reasoning, autonomous code generation, and iterative self-correction, future DREAM Challenges may incorporate agent leaderboards as a standard track alongside human competitors, creating a dynamic benchmark for the state of autonomous scientific discovery. Finally, the Challenge infrastructure, evaluation pipelines, and model-to-data framework developed here can be re-used and adapted for future challenges targeting other (neurodegenerative) diseases including Parkinson’s disease, frontotemporal dementia, and Lewy body disease, for which comparable single-cell atlases are actively being developed.

## Acknowledgements

We thank all challenge participants for their contributions, scientific creativity, and engagement throughout the open and final phases of the SEA-AD DREAM Challenge (the list can be found in Supplementary Table 3 and also here: https://www.synapse.org/Team:3542566). The breadth and quality of submitted solutions made this community benchmark possible.

The SEA-AD DREAM Challenge was supported by the National Institute on Aging (NIA) grants U24AG061340 supporting the AD Knowledge Portal Data Coordination Center, and U19AG060909 to the Seattle Alzheimer’s Disease Brain Cell Atlas (SEA-AD) consortium. This study was also supported by the March of Dimes Prematurity Research Center at UCSF (TO and MS). SEA-AD data were generated from postmortem brain tissue donated to the University of Washington BioRepository and Integrated Neuropathology (BRaIN) laboratory and Precision Neuropathology Core, which is supported by the UW Alzheimer’s Disease Research Center (NIA P30AG066509, previously P50AG005136), the Adult Changes in Thought (ACT) Study (NIA U19AG066567). ACT Data collection was additionally supported, in part, by prior funding from NIA (U01AG006781), and the Nancy and Buster Alvord Endowment. We thank the participants of the ADRC and the ACT study for the data they have provided and the many ADRC and ACT investigators and staff who steward that data. You can learn more about UW ADRC at https://depts.washington.edu/mbwc/adrc and ACT at https://actagingstudy.org/. We thank AMP-AD, Sage Bionetworks, and the Open Data Registry on AWS for hosting various SEA-AD datasets.

## Author contributions

H.L., G.A., L.F., L.H.,T.O., M.S., G.S., K.J.T., J.Z., M.I.G. designed the challenge. J.S.R. and L.A. conceived the AI Agent Track. G.A., L.F., L.H. created infrastructures for the challenge. H.L., E.S.K., K.J.T., M.I.G. organized the data for the challenge. N.K., V.C. performed the benchmarking analysis. H.L., N.K., G.A., wrote the majority of the manuscript. D.A., L.C., E.C., I.G.V., G.G., L.J.H., X.L., E.L., P.L., I.N., F.O., T.O.Y., I.Ri., I.Ro. participated in the challenge and contributed to the manuscript writing. G.A., L.F., L.H.,T.O., M.S., G.S., K.J.T., J.Z., M.I.G. supervised the challenge and the manuscript writing.

## Declaration of interests

J.S.R. reports in the last 3 years funding from GSK and Pfizer & fees/honoraria from Travere Therapeutics, Stadapharm, Astex Pharmaceuticals, Owkin, Pfizer, Vera Therapeutics, Grunenthal, Tempus and Moderna.

## STAR Methods

### Key resources table

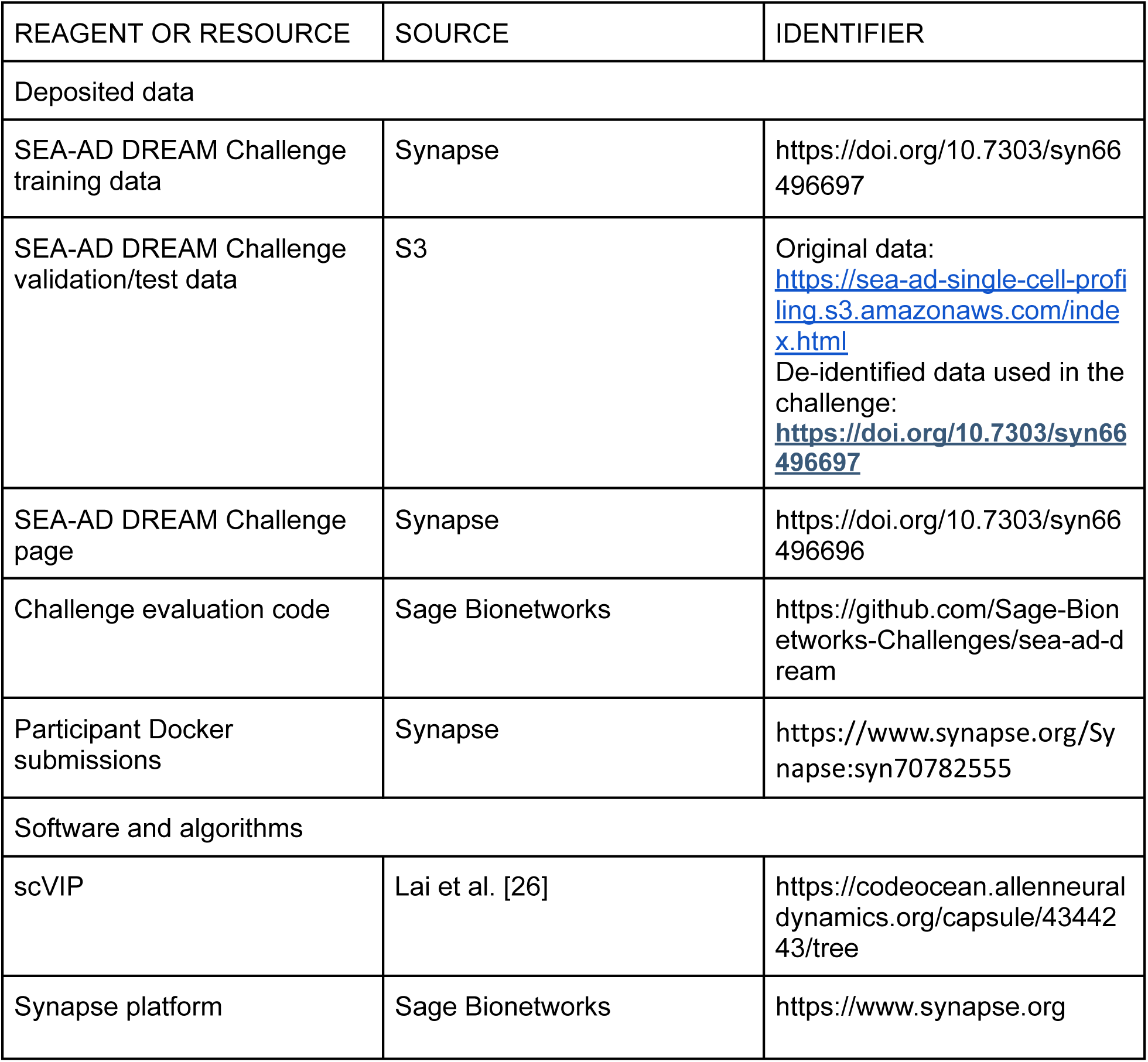

### Resource availability

#### Lead contact

Mariano Gabitto

#### Materials availability

These challenges did not generate new materials.

#### Data and code availability

- The challenge data has been deposited at Synapse and is publicly available as of the date of publication at https://doi.org/10.7303/syn66496697 and at https://sea-ad-single-cell-profiling.s3.amazonaws.com/index.html
- All original code has been deposited at Synapse and is publicly available as of the date of publication at https://github.com/Sage-Bionetworks-Challenges/sea-ad-dream and at https://www.synapse.org/Synapse:syn70782555.
- Any additional information required to reanalyze the data reported in this paper is available from the lead contact upon request.

### Experimental model and study participant details

#### Training data and validation/test data cohort

The training cohort comprised 84 donors recruited through the ACT study, representing the full spectrum of AD neuropathological severity (Supplementary Figure 2). All donors have accompanying clinical and demographic metadata (donor ID, sex, age at death, APOE4 genotype, years of education, post-mortem interval, race, and Hispanic/Latino ethnicity) as well as complete neuropathological staging (ADNC, Braak, Thal, CERAD, LATE-NC, Lewy Body Disease stage) and quantitative immunohistochemical burden scores (percent positive area for 6E10, AT8, NeuN, GFAP, pTDP43, and aSyn) available in AnnData.obs.

The validation/testing cohort consisted of 42 donors from the 84 donors in the training cohort. The donor demographics on this subset can be found in Supplementary Figure 3.

### Methods details

#### Training and validation/testing data acquirement

Training data were generated from two brain regions: the middle temporal gyrus (MTG, Brodmann area 21) and the dorsal frontal cortex (DFC, Brodmann area 9). Single-nucleus RNA-seq libraries were prepared using 3’ 10x Genomics Chromium v3.1 or 10x Multiome protocols at the Allen Institute for Brain Science, following a common standardized pipeline across all donors. Raw reads were de-multiplexed and mapped to the official 10x Genomics 2020-A human reference index (GRCh38), yielding expression measurements for 36,601 features per cell. Data are provided as AnnData (.h5ad) objects with log-normalized counts per 10,000 (adata.X) and raw UMI counts (adata.layers[’UMIs’]) for each nucleus. Cell type annotations follow a three-level hierarchical taxonomy comprising 3 Classes, 24 Subclasses, and 139 Supertypes, as described in the SEA-AD reference atlas.

Quantitative immunohistochemical (IHC) pathology measures were obtained from whole-slide images of tissue sections immediately adjacent to those used for snRNA-seq. IHC staining was performed by the University of Washington Biorepository and Integrated Neuropathology (BRaIN) Laboratory using antibodies targeting the following proteins: 6E10 (amyloid-beta, pan-plaque), AT8 (phospho-tau Ser202/Thr205), NeuN (RBFOX3, neuronal marker), GFAP (astrocytic marker), pTDP43 (phosphorylated TDP-43), and aSyn (alpha-synuclein, Lewy body marker). Digital image analysis was performed using HALO v.3.4.2986 (Indica Labs, Albuquerque, NM). For each marker, the primary quantitative output was the percent of voxels staining positive in the grey matter of the profiled cortical region. Representative staining and HALO-derived quantification masks are shown in Supplementary Figure 4. Pathological protein burden (6E10, AT8, pTDP43, aSyn) and cellular marker burden (NeuN, GFAP) across cortical layers are also visualized as donor-ordered heatmaps in Supplementary Figure 4, with values normalized to z-scores and adjusted by a moving average along the cognitive-pathological spectrum (CPS) [35].

Validation and test data used for leaderboard scoring and final round scoring were generated from the superior temporal gyrus (STG) and inferior temporal gyrus (ITG) [31]. The datasets were processed and annotated using the same pipeline as the training data and provided in identical AnnData (.h5ad) format, with transcriptomic measurements but without neuropathological or immunohistochemical labels accessible to participants. The use of a distinct brain region for validation and testing ensures that high-performing models demonstrate generalization across cortical areas rather than region-specific signal memorization.

#### DREAM challenge powered by Sage Bionetworks

##### Overall challenge structure

An overview of the Challenge is shown in Figure 6. All Challenge elements were supported by the Synapse collaborative research platform (https://www.synapse.org; Sage Bionetworks, Seattle, WA), including documentation, access to training data, model submission, automated evaluation, leaderboards, and participant discussion forums. To gain access to training data, teams were required to register on Synapse and accept a Data Use Agreement (DUA) restricting use of SEA-AD data to Challenge purposes and requiring compliance with the ethical guidelines established by the SEA-AD data access policy. The Challenge comprised two sequential prediction Tasks (Task 1: categorical neuropathological staging; Task 2: continuous quantitative pathological burden), each evaluated independently with dedicated leaderboards and primary metrics (see Quantification and Statistical Analysis).

**Figure 6:**
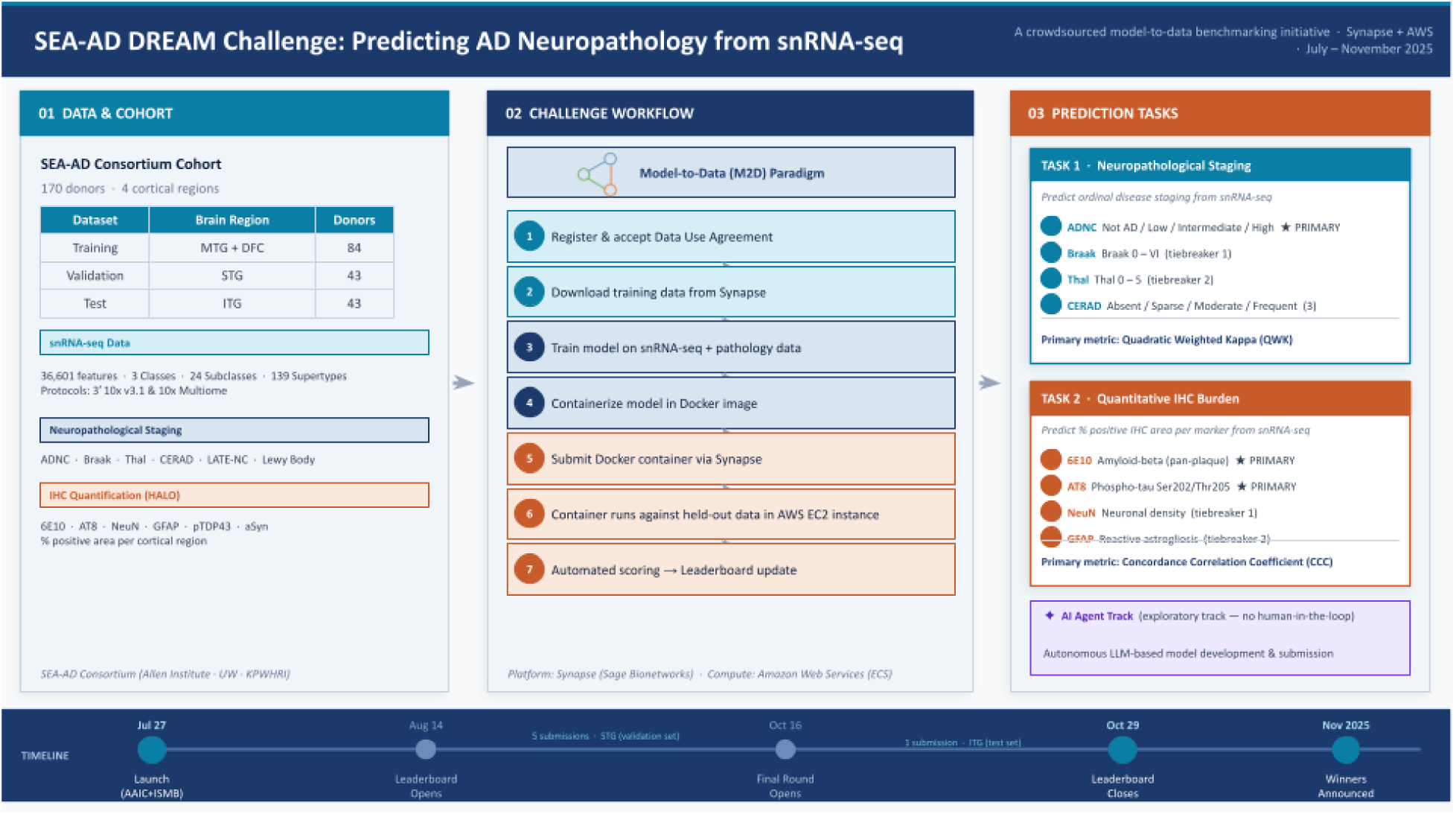
Challenge Overview. Panel 01 — Data & Cohort (teal header): cohort summary table (Training MTG+DFC 84, Validation STG 43, Test ITG 43), plus colour-coded blocks for snRNA-seq specs (36,601 features, 3 Classes / 24 Subclasses / 139 Supertypes), neuropathological staging variables, and IHC markers with HALO note. Panel 02 — Challenge Workflow (navy header): the model-to-data paradigm callout, followed by a 7-step numbered flow (register → download → train → dockerize → submit → evaluation on cloud compute → leaderboard update), with colour-coded step groups (teal = participant-side, navy = model, orange = platform). Panel 03 — Prediction Tasks (orange header): Task 1 (Neuropathological Staging) with all four targets (ADNC ★ PRIMARY → Braak → Thal → CERAD) and QWK metric; Task 2 (Quantitative IHC Burden) with 6E10 + AT8 ★ PRIMARY → NeuN → GFAP and CCC metric; plus the purple AI Agent Track badge. Timeline strip (navy): seven milestones from Jul 27 launch through Nov 2025 winner announcement, with phase labels showing submission limits and dataset used.

The Challenge was implemented as a model-to-data competition. Rather than distributing the held-out validation cohort to participants, teams were required to encapsulate their trained models and all software dependencies within Docker containers. Upon submission through the Synapse platform, containers were transferred to the evaluation infrastructure and executed against the validation data in a secure, isolated environment; raw donor-level validation data did not leave the controlled compute environment at any point. This architecture prevents post-hoc model tuning to the test set and ensures that reported performance metrics reflect genuine out-of-sample generalization.

Docker container evaluation was performed on Amazon Web Services (AWS) using Elastic Compute Cloud (EC2) instances. Container execution was orchestrated through Synapse evaluation queues and automated workflow scripts maintained by the Sage Bionetworks challenge infrastructure team. Participants had no direct access to the evaluation compute environment or to validation labels. All containers were executed under standardized resource constraints (110 GB of system memory, 2 GB of shared memory, 1 vCPU, and 3-hour runtime limit) to ensure equitable evaluation. To maintain data privacy and prevent models from accessing external information, all network access were disabled. Prediction output files generated by each container were automatically captured and scored against held-out ground-truth labels using the primary and tiebreaker metrics described below.

The open (leaderboard) phase ran from August 4 to October 29, 2025 (Final round from Oct 16-29). During this phase, teams received real-time leaderboard feedback on Task 1 and Task 2 performance and were permitted a maximum of 5 total scored submissions across both Tasks, with the top-performing submission per Task designated as the team’s official final submission at the close of the open phase. Final rankings were computed from the official submissions on the same held-out validation data. To be included as participants in this publication, teams were required to: (i) make their submission code publicly available under an open-source license; (ii) provide a written methods summary; and (iii) complete a post-challenge survey documenting modeling strategies, key features, and insights. This eligibility requirement was communicated to all registered teams at the time of Challenge registration.

##### Participant engagement

Information about the Challenge was disseminated through the DREAM Challenges website (https://dreamchallenges.org), Sage Bionetworks communications channels, and broader scientific mailing lists. Announcements were also shared through social media platforms including LinkedIn, and through direct outreach to relevant research communities in computational biology, machine learning, and Alzheimer’s disease research. To lower the barrier to participation, Challenge organizers prepared a detailed participant guide, an example Docker container with a functional example model, and standardized input data format documentation, all hosted on the Synapse project page.

Organizers held virtual webinars at Challenge launch and at key milestones throughout the open phase to describe the SEA-AD dataset, explain the Task definitions and evaluation metrics, and answer participant questions. Ongoing engagement was maintained through the Synapse discussion forum, where organizers responded to technical and scientific questions in near real-time. In order to preserve model environments for portability and reproducibility, participants were required to submit fully self-contained Docker environments including all programming language dependencies, trained model weights, and inference code needed to generate predictions on a standardized input .h5ad file.

#### Top-performing teams’ models in the human track

The three top-performing teams - consistent across both tasks - describe their approaches below (the order of the teams is alphabetical):

#### Top performing team: Team BioICAR

Team BioICAR developed a two-stage modeling framework that first generated predictions from each cell supertype and then integrated these predictions to produce donor-level outputs.

Rather than modeling individual cells directly, the method summarized each donor–supertype pair by averaging gene expression across cells of the same supertype. These donor–supertype profiles were then jointly compressed using principal component analysis across all supertypes, with the top 2,048 principal components retaining approximately 95.3% of the explained variance.

In the first stage, separate models were trained for each cell supertype. For Task 1, BioICAR used L2-regularized multinomial logistic regression to estimate the probability of each ADNC class. For Task 2, they used L2-regularized ridge regression to predict quantitative pathological burden for each marker, 6E10 and AT8. This cell-supertype-specific modeling strategy was motivated by the instability of direct gene–supertype feature selection across cross-validation folds and by the need to reduce the dimensionality of the donor-level feature space.

In the second stage, the cell-supertype-level predictions were concatenated and used as input to a final donor-level model. For Task 2, donor metadata, including sex, race, age at death, years of education, post-mortem interval, and APOE genotype, were also included at this stage. The final model was selected based on validation performance: ridge regression was used for AT8, whereas XGBoost was used for 6E10 to capture potential nonlinear associations. Five-fold cross-validation was used throughout to tune regularization parameters for linear models and hyperparameters for XGBoost.

Although the framework relied on aggregated donor–supertype profiles, it retained a degree of interpretability. Because the first-stage models were linear, predictive features could be examined through model coefficients and mapped back to gene-level contributions through PCA back-projection.

#### Top performing team: Team CMC-TJU

In contrast to BioICAR, Team CMC-TJU used a more restricted feature set and trained predictors at the level of broad cellular classes rather than individual cell supertypes. The method limited the input space to 3,430 nervous system genes, of which 3,157 were included in the training data made available to the participants. These genes are enriched for primate- and human-specific sequence motifs (“pyknons”) [29], and grouped cells into three broad classes: GABAergic neurons, glutamatergic neurons, and non-neuronal cells. To increase the number of training observations, cells within each donor–class pair were divided into 100 equal subgroups, each summarized by mean expression across the pyknon-associated genes. Donor metadata, including sex, years of education, and APOE genotype, were appended to each subgroup-level expression profile, with categorical variables one-hot encoded.

Separate LASSO regression models were trained for each outcome variable using glmnet with α = 1 and data from both the MTG and DFC regions. The regularization parameter λ was selected by cross-validation using the training data only, and final models were fit using all available training data. Model performance was estimated using an 80/20 donor-level train/test split, with no donor overlap between splits.

At inference, the same preprocessing and regularization parameters were applied to generate multiple subgroup-level predictions for each donor. These predictions were aggregated into donor-level outputs using majority voting for Task 1 ADNC class prediction and averaging for Task 2 prediction of 6E10 and AT8 burden. Task 2 predictions outside the range of 0 to 100 were clipped to the nearest boundary.

#### Top performing team: Team gisl7

For task 1, team gisl7 performed 3-fold cross-validation across a range of machine-learning models implemented in Scikit-learn to assess the relevance of the model features heuristically derived from the donor level metadata and single-cell RNA-Seq data.

The machine learning models include CatBoost, Random Forest, Extra Trees, Gradient Boosting, HistGradientBoosting, XGBRF, LightGBM, AdaBoost, Ridge regression, Lasso regression, Elastic Net, Support Vector Regression, and K-Nearest Neighbors. For each neuropathological outcome, the optimal model and the mapping from numerical predictions to ordinal outcomes were selected based on the quadratic weighted kappa (QWK) metric. The selected model was subsequently retrained using the full training dataset.

Based on diagnostic analyses conducted during model development, donor-level metadata were selected as the model features, including sex, age at death, years of education, post-mortem interval, and APOE genotype. One-hot encoding was applied to sex and APOE genotype.

During model evaluation, team gisl7 observed that models trained exclusively on metadata achieved unexpectedly high, and in some cases perfect, leaderboard performance. Because this pattern could affect the reliable identification of meaningful predictive features, the observation was reported to the challenge organizers as a potential data leakage or evaluation-design concern. Given these findings, a metadata-based model was submitted as the final model.

For task 2, team gisl7 summarized each donor and brain region (MTG and A9) using gene-level expression fractions, defined as the fraction of cells with nonzero expression for each gene.

These donor-by-gene expression-fraction matrices formed the basis for downstream feature selection and model training.

To select robust expression features, Team gisl7 ranked genes by their association with each pathology outcome using mutual information [36] estimated with a spline-based estimator [37] implemented in the cafr R package. Mutual information was used as a non-parametric measure of association between donor-level gene-expression fractions and pathology burden. The estimates were normalized so that the maximum possible value was one, and the corresponding Pearson correlation was used to determine the direction of association, with values clipped to zero when the Pearson correlation indicated a negative association. Positive and negative features were then selected separately to prioritize genes with robust and directionally interpretable associations with the pathology outcomes.

To reduce sensitivity to sampling variation in cellular composition, the team applied this ranking procedure within a stratified subsampling framework. For each brain region, cells were stratified into 20 non-overlapping subsets with similar size and supertype composition. For each subset, the team generated a donor-by-gene expression-fraction matrix, ranked genes by mutual information with 6E10 or AT8 burden, and aggregated the resulting rankings using the median rank.

For each brain region and target, the team retained the top-ranked positively and negatively associated genes, including features selected from the full donor set and from donors above predefined high-burden thresholds, defined as 6E10 > 6 or AT8 > 4. This yielded 20 mutual-information-selected expression features per target. For 6E10 prediction in MTG, where the initially selected features showed lower predictive performance, the team further filtered candidate genes based on cross-validated model performance and retained the top 10.

In addition to single-gene expression features, Team gisl7 constructed gene-pair co-expression features from the microglia–perivascular macrophage subclass, selected because of its relevance to Alzheimer’s disease pathology and consistent representation across brain regions [15,38]. Genes expressed in fewer than 10% of cells were removed, and the 1,000 most variable genes were retained, yielding 499,500 candidate gene pairs. For each donor, pairwise co-expression was estimated from single-cell profiles using the Ledoit–Wolf shrinkage estimator to improve covariance stability.

To reduce noise in the co-expression estimates, cells from each donor were divided into four stratified groups. Pairwise correlations were computed within each group, and the median correlation across groups was used as the stabilized donor-level co-expression value. Robust gene pairs were then selected through repeated split-sample validation. Across 10 iterations, donors were divided into discovery and replication sets; candidate gene pairs were identified by Spearman correlation with the pathology target in the discovery set and retained only if they showed consistent direction and comparable correlation strength in the replication set. The final consensus set consisted of reproducible gene pairs that appeared across multiple iterations and contributed to model performance.

For model training, Team gisl7 combined mutual-information-selected expression fractions, stabilized gene-pair co-expression features, and donor-level covariates, including sex, APOE genotype, age at death, years of education, and post-mortem interval. Supervised regression models were trained separately for each pathology target. To avoid donor-level leakage while maintaining balanced target distributions, the team used stratified group K-fold cross-validation, grouping samples by donor ID and stratifying pathology values by quantile bins. Model performance was evaluated using out-of-fold predictions and quantified by concordance correlation coefficient.

The team evaluated several regression algorithms, including gradient boosting regression with Huber loss, extremely randomized trees, and random forest regression. The best-performing model for each target was selected based on cross-validated out-of-fold concordance correlation coefficient. To assess cross-region generalization, models for 6E10 were trained on MTG, where the amyloid-β burden showed a broader signal range, and evaluated on A9; conversely, models for AT8 were trained on A9 and evaluated on MTG. The final submitted predictions were generated from the best-performing cross-region models.

#### Complementary organizer-side model

To apply scVIP to the DREAM Challenge, we followed the training procedure previously used for the combined SEA-AD and ROSMAP analysis [26]. We used a semi-supervised training strategy in which all staging and quantitative neuropathology measurements for the held-out STG and ITG datasets were marked as missing. Because donor overlap and batch relationships between the training and held-out regions were not provided to the model, each STG and ITG donor was assigned a new donor identifier and batch identifier. The training dataset included ROSMAP dorsolateral prefrontal cortex (DLPFC) glial data together with SEA-AD middle temporal gyrus (MTG) and dorsal frontal cortex (DFC/A9) glial data, as in the previous scVIP analysis, and the held-out SEA-AD STG and ITG datasets were included without observed pathology labels for semi-supervised inference. Together, these datasets comprised 1,146,585 glial cells. Because neuropathological measurements differ across cohorts, we used shared staging variables, including Braak stage, Thal phase, CERAD score, and ADNC, as common supervision across datasets, while allowing cohort-specific quantitative measurements to be partially observed. These cohort-specific measures included percent 6E10 and AT8 burden for SEA-AD, and diffuse plaque density, neuritic plaque density, amyloid density, and neurofibrillary tangle density for ROSMAP. All the neuropathology measurements are log-transformed just as in [26]. We first trained scVI on the full combined dataset to learn cell-level embeddings, using cohort as the batch key and batch identifier as an additional covariate. scVIP was then initialized from the trained scVI model and further optimized to refine the cell-level embeddings while learning donor-level representations and neuropathology prediction parameters. The optimization parameters are the same as in [26].

#### Models in the Agent track

Submissions to the Agent Track followed the same Docker-based evaluation pipeline, compute constraints, and scoring metrics as the main track. Each team additionally submitted the agent’s thought-process log and tool-use trace, and each human collaborator completed an AI-contribution checklist reporting involvement level (A: human-generated; B: mostly human, AI-assisted; C: mostly AI, human-assisted; D: AI-generated) across four categories: hypothesis development, experimental design and implementation, data analysis and interpretation, and observed AI limitations. Two teams submitted. The first (Nguyen & Zhan, New Mexico Tech) used a single LLM under human supervision for code synthesis and debugging (self-rated C across all categories, with the authors noting that the submission was a human-controlled execution of LLM-generated code rather than an autonomous controller); the final pipeline used 3,000-component PCA, per-endpoint MLP classifiers for ordinal targets, and Random Forest regressors for continuous targets, with donor-level aggregation. The second (UWisc-Madison) implemented a multi-agent AutoML framework comprising an Agent Manager (coordinator), a Prompt Agent (instruction parsing), Data and Model Agents (Gemini 2.5 Pro, Google) for domain-aware planning, and an Operation Agent (Claude Code 4.5, Anthropic) for code synthesis, execution, and self-debugging, organized as multi-round plan → code → validate → refine loops; the final pipeline used 20-component PCA per cell-type Class over 2,000 HVGs intersected across A9 and MTG, donor-level aggregation with metadata, and an ordinal XGBoost ensemble (three cumulative binary classifiers) for ADNC with multiclass XGBoost for Braak, Thal, and CERAD.

### Quantification and statistical analysis

#### Benchmarking

##### Submission Format and Validation

Each submission was required to provide one prediction per donor, formatted as a CSV file indexed by Donor ID. For Task 1, submissions predicted four neuropathological staging scores: ADNC, Braak, CERAD, and Thal. For Task 2, submissions predicted the percentage of positively stained area for four immunohistochemical markers: 6e10, AT8, NeuN, and GFAP. Predictions for two additional neuropathological features (LATE staging and Highest Lewy Body Disease for Task 1; α-synuclein and pTDP-43 for Task 2) were accepted as optional columns.

Upon receipt, each submission was automatically validated to confirm the presence of all required columns, the absence of missing or duplicate Donor IDs, that categorical values belonged to their respective accepted label sets, and that continuous values fell within the valid range. Submissions failing any of these checks were returned with an error message and excluded from scoring.

##### Scoring Metrics

Task 1 submissions were evaluated using the Quadratic Weighted Kappa (QWK), which extends Cohen’s kappa by applying quadratic weights to penalize predictions proportionally to their ordinal distance from the ground truth. QWK was computed for each of the four predicted staging scores by first mapping their ordinal labels to integers according to disease progression order. The primary ranking metric was ADNC QWK, with Braak, Thal, and CERAD QWK used sequentially as tiebreakers. Mean absolute error (MAE) and Spearman’s rank correlation were computed as secondary, non-ranking diagnostics.

Task 2 submissions were evaluated using Concordance Correlation Coefficient (CCC), which jointly captures both the Pearson correlation and the agreement between predicted and observed means and variances. CCC was computed independently for each of the four immunohistochemical markers. Final ranking was determined by the average CCC across the 6e10 and AT8 markers, which represent amyloid-β and phosphorylated tau pathology respectively, with NeuN and GFAP CCC used as tiebreakers. Mean squared error (MSE) and Pearson R² were computed as secondary diagnostics.

##### Top Performer Determination

To identify statistically distinguishable top-performing submissions, we applied a bootstrapping procedure combined with a Bayes Factor (BF) analysis. All Final Round prediction files were merged with the ground truth into a single combined data frame, and the scoring pipeline was rerun on the original (non-resampled) data as a pre-bootstrapping verification step to confirm that computed scores matched the published leaderboard values exactly. The core procedure then drew N=10,000 bootstrap samples with replacement from the donor-level predictions, re-scoring each submission on every resample using the primary challenge metric (QWK for Task 1; average CCC across 6e10 and AT8 for Task 2). The BF for each submission was computed relative to the top-performing reference submission (BF=0) by taking the ratio of bootstrap iterations in which a given method outperformed (or matched) the reference versus those in which it did not. A BF≤3 was used as the threshold for declaring statistical equivalence with the top performer, following established DREAM Challenge conventions; submissions meeting this criterion were co-declared Top Performers alongside the reference method.

#### Data leakage analysis

We performed three tests. In Test 1, metadata and labels were jointly shuffled across donors; under this setting, donors effectively “exchange cell populations”, resulting in models relying primarily on metadata should retain their performance. In Test 2, only metadata were shuffled while labels were preserved; under this setting, donors “exchange metadata profiles”, models that do not depend on metadata should remain largely unaffected. In Test 3, the order of cells was shuffled to assess whether any methods implicitly relied on donor or cell ordering. All teams were evaluated by comparing their official leaderboard performance with their performance under these perturbation settings. To estimate variability in the top-performing methods, the analyses were repeated 10 times for the top three teams on each leaderboard. The same evaluation framework was then applied to Task 2 submissions.

#### Feature importance analysis

For BioICAR, we interpreted the two-stage framework by tracing feature contributions through both levels of the model. The second-stage donor-level model was used to identify informative cell supertypes, because each input block represented the prediction generated from one supertype-specific model. Coefficients from linear models or feature-importance estimates from tree-based models therefore provided supertype-level importance scores for the final donor-level prediction. To relate these supertype-level contributions back to gene expression, we then examined the first-stage supertype-specific models. Because these models were trained on PCA-compressed donor–supertype expression profiles, model weights were mapped from principal-component space back to the original gene space using PCA back-projection, providing an approximate gene-level interpretation of the expression programs driving each informative supertype-level predictor.

For CMC-TJU, feature interpretation relied on the sparsity of the glmnet models used for both Task 1 and Task 2. We extracted model coefficients in R to identify predictors retained by the fitted models and to examine whether each retained predictor was positively or negatively weighted.

For gisl7, feature importance was assessed from the tree-based regression models used for Task 2 by TreeSHAP on the final tree models, grouped by attribute types. These estimates were used to prioritize expression-fraction features, microglia–perivascular macrophage co-expression features, and donor metadata variables associated with the final predictions.

## Supplementary Figures

**Supplementary Figure 1:**
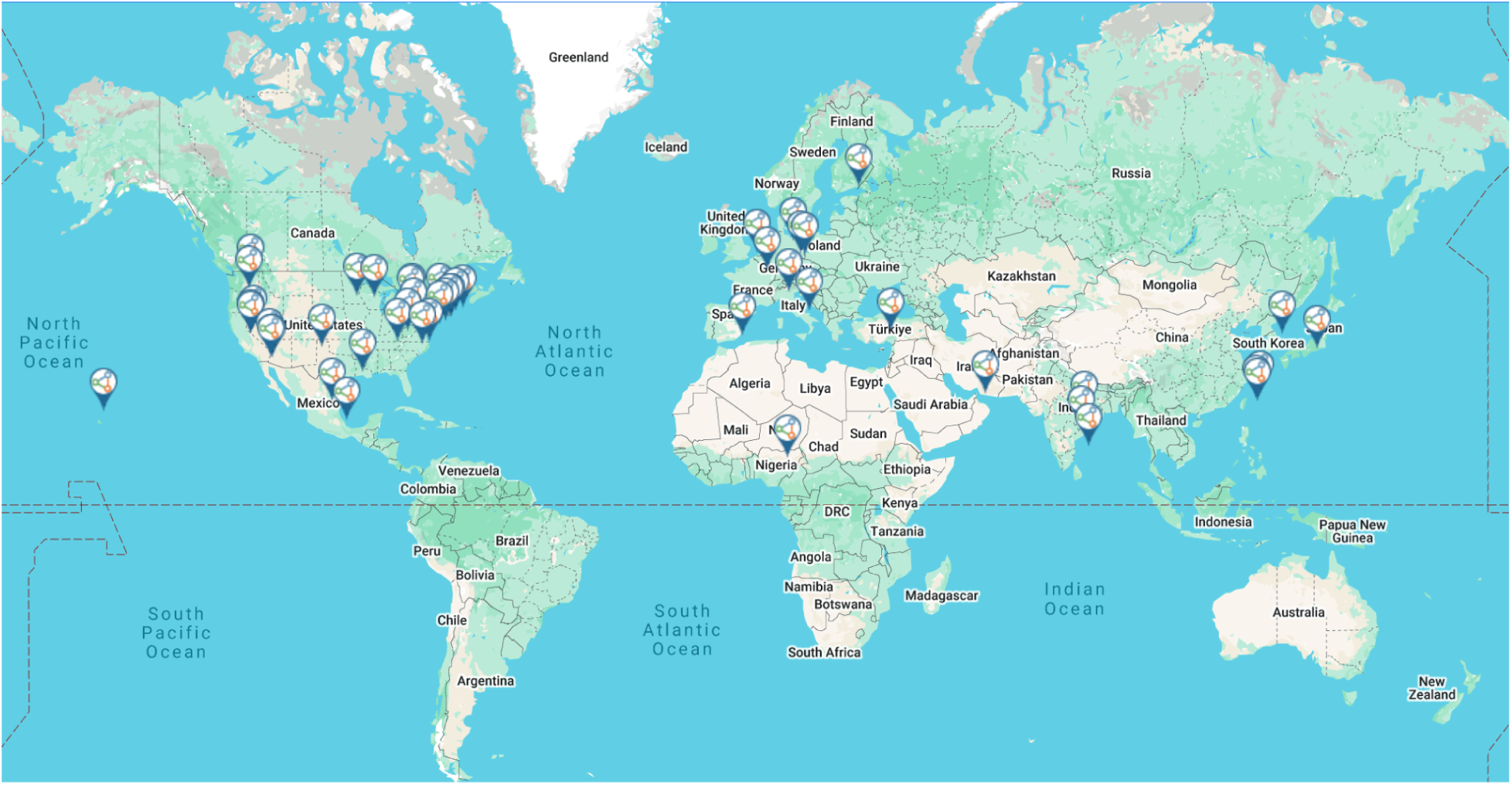
Distribution of the participating individuals/teams

**Supplementary Figure 2:**
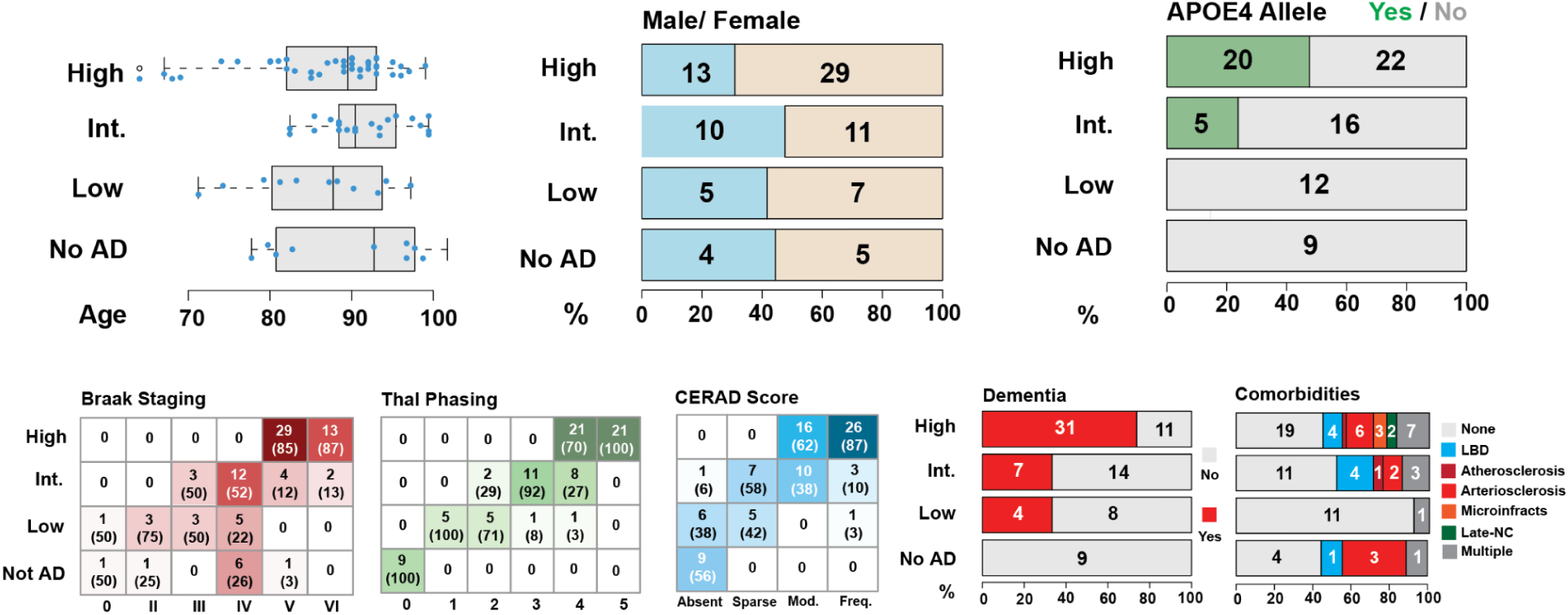
Training datasets demographics: (top) SEA-AD cohort demographics, depicting age at death, biological sex and APOE4 allele, and dementia status stratified according to Overall AD Neuropathological Change (ADNC). Age at death is represented by box-and-whisker plots with the box representing the interquartile range (IQR) and the whiskers representing 1.5 times the IQR. The solid line indicates the median. SEA-AD cohort composition stratified according to ADNC versus Braak stage (measure of pTau tangle distribution), Thal phase (measure of amyloid plaque distribution), and CERAD score (measure of neuritic plaque burden) as heatmaps, with dementia and comorbidities as bar plots. The number of donors in each box and the fraction are shown in parentheses.

**Supplementary Figure 3:**
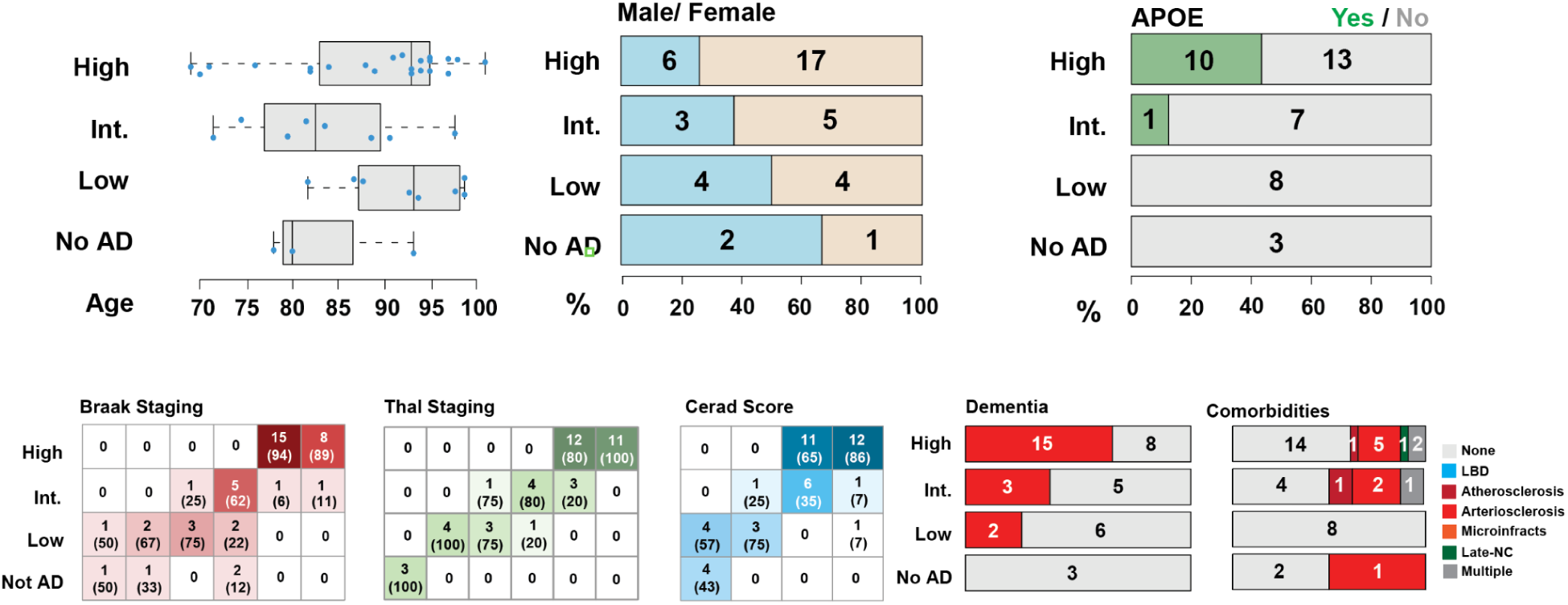
Validation and test datasets demographics: (top) SEA-AD cohort demographics, depicting age at death, biological sex and APOE4 allele, and dementia status stratified according to Overall AD Neuropathological Change (ADNC). Age at death is represented by box-and-whisker plots with the box representing the interquartile range (IQR) and the whiskers representing 1.5 times the IQR. The solid line indicates the median. SEA-AD cohort composition stratified according to ADNC versus Braak stage (measure of pTau tangle distribution), Thal phase (measure of amyloid plaque distribution), and CERAD score (measure of neuritic plaque burden) as heatmaps, with dementia and comorbidities as bar plots. The number of donors in each box and the fraction are shown in parentheses.

**Supplementary Figure 4:**
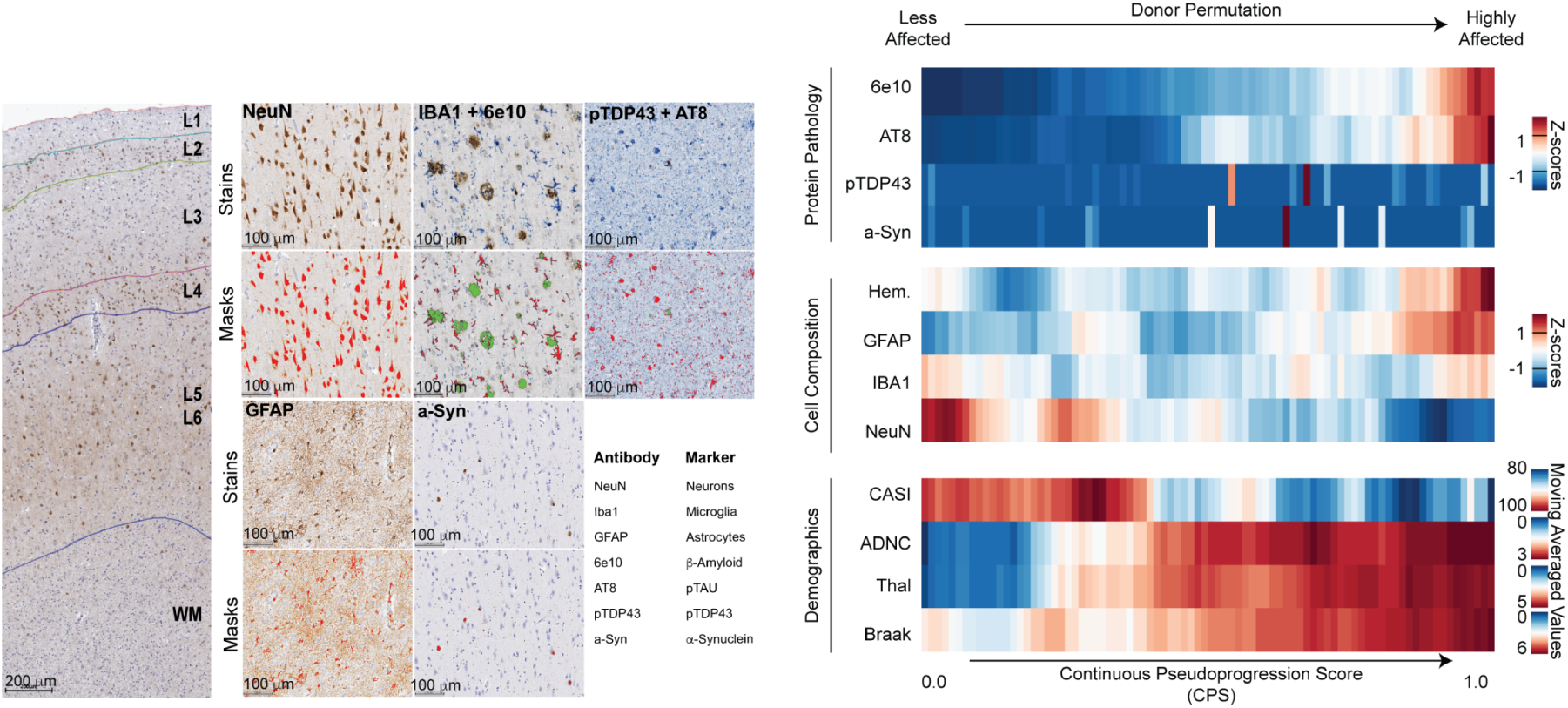
(Left) Representative cortical column visualized with immunohistochemistry (IHC). Cortical layers (L1–L6) and white matter are indicated. Immunostaining was performed in the entire SEA-AD cohort (n = 84). Higher-powered micrographs showing IHC staining for protein aggregates and cellular populations. Bottom, masks showing positive voxels generated by HALO in red for single staining and both red and green for duplex staining. (Right, top) Heatmap showing the number of pathological protein objects detected per unit area across all cortical layers in each donor, ordered along a CPS. All values were converted to z-scores and adjusted according to a moving average. (Right, middle), Heatmap showing the number of cellular objects detected per unit area across all cortical layers, ordered along the CPS. Hem, hematoxylin+ nuclei; GFAP, IBA1 and NeuN indicates the number of positive cells. All values were converted to z-scores and adjusted according to a moving average. (Right, bottom), Heatmap showing the cognitive scores at the last visit (CASI) and AD pathology stage (ADNC, Thal, Braak), ordered along the CPS. All values were adjusted according to a moving average.

## Supplementary Tables

**Supplementary Table 1.1.**
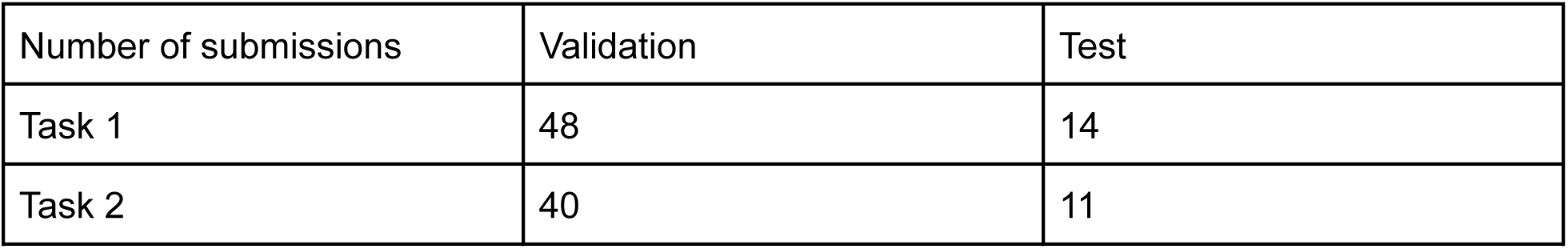
Number of scored submissions in the validation round (i.e., Leaderboard Round) and in the test round (i.e., Final Round) for Task 1 and 2.

**Supplementary Table 1.2.** Number of teams that participated in the validation round and in the test round for Task 1 and 2.

| Number of teams | Validation | Test |
| --- | --- | --- |
| Task 1 | 17 | 14 |
| Task 2 | 15 | 11 |

## Notes

https://doi.org/10.7303/syn66496696

### Competing Interest Statement

The authors have declared no competing interest.

### Summary of Updates

Including more features identified by the top three DREAM challenge participating teams.

